# GABA excitatory actions in cerebrospinal-fluid contacting neurones of adult mouse spinal cord

**DOI:** 10.1101/2022.12.14.520067

**Authors:** Priscille Riondel, Nina Jurčić, Jérôme Trouslard, Nicolas Wanaverbecq, Riad Seddik

**Author notes:** Correspondence should be addressed to (R. SEDDIK) or (N. WANAVERBECQ). RS and NW contributed equally to this work as Principal Investigators (PIs).

## Abstract

Spinal cerebrospinal fluid-contacting neurons (CSF-cNs) form an evolutionary conserved bipolar cells population localized around the central canal of all vertebrates. CSF-cNs were shown to express molecular markers of neuronal immaturity into adulthood, however the functional relevance of their incomplete maturation remains unknown. Neuronal maturation is classically associated with the expression of the K^+^-Cl^-^ cotransporter 2 (KCC2), allowing chloride (Cl^-^) extrusion and hyperpolarising GABA transmission. Here, we show no detectable expression of KCC2 in CSF-cNs of adult mouse spinal cord. Accordingly, lack of KCC2 expression results in low Cl^-^ extrusion capacity in CSF-cNs under high Cl^-^ load in whole-cell patch-clamp. Using cell-attached recordings, we found that activation of ionotropic GABA_A_ receptors induced a dominant depolarising effect in 70% of CSF-cNs recorded with intact intracellular chloride concentration. Moreover, in these cells, depolarising GABA-responses can drive action potentials as well as intracellular calcium elevations by activating voltage-gated calcium channels. CSF-cNs express the Na^+^-K^+^-Cl^-^ cotransporter 1 (NKCC1) involved in Cl^-^ uptake and its inhibition by bumetanide blocked the GABA-induced calcium transients in CSF-cNs. Finally, we show that activation of metabotropic GABA_B_ receptors did not mediate hyperpolarisation in spinal CSF-cNs, presumably due to the lack of expression of G protein-coupled inwardly rectifying potassium (GIRK) channels. Together, these findings outline CSF-cNs as a unique neuronal population in adult spinal cord with immature Cl^-^ homeostasis and no hyperpolarising GABAergic signalling but rather generation of excitation and intracellular calcium modulation. GABA may therefore promote CSF-cNs maturation and integration into the existing spinal circuit.

**Key points:**

- CSF contacting neurones (CSF-cNs) are located around the central canal of spinal cord across all vertebrates.
- CSF-cNs express canonical markers of immature neurons during adulthood in mice but the impact of such persistent immaturity on their chloride (Cl^-^) homeostasis as well as GABAergic signalling were not addressed yet.
- Here, we show that spinal CSF-cNs express the Na^+^-K^+^-Cl^-^ cotransporter 1 (NKCC1) involved in Cl^-^ uptake but not the K^+^-Cl^-^ cotransporter 2 (KCC2) classically allowing Cl^-^ extrusion.
- As a result of intracellular Cl^-^ accumulation, GABA does not mediate inhibition in most CSF-cNs but rather excitation and intracellular Ca^2+^ elevations through the activation of voltage-gated Ca^2+^ channels.
- Excitatory GABAergic signalling associated with intracellular calcium modulation may underlie the maturation and integration of CSF-cNs into the spinal circuit of adult mice.

## Introduction

Chloride-permeable ionotropic GABA_A_ receptors (GABA_A_-Rs) mediate inhibition in adult brain, by generating an outward chloride (Cl^-^) current and hence a hyperpolarisation of the membrane potential. However, under conditions of intracellular Cl^-^ accumulation, GABA_A-_Rs were shown to promote inward Cl^-^ current and depolarising responses (Owens & Kriegstein, 2002). The intracellular Cl^-^ concentration in neurones is mainly determined by two Cl^-^ transporters, the K^+^-Cl^-^ cotransporter 2 (KCC2) and the Na^+^-K^+^-Cl^-^ cotransporter 1 (NKCC1), whose expression/efficacy is developmentally regulated (Payne *et al*., 2003). In neuronal precursors and immature neurones, NKCC1 predominates and cotransports Cl^-^ ion into cells using the Na^+^ gradient, thus allowing an increase of intracellular Cl^-^ concentration and depolarising action of GABA. During neuronal maturation NKCC1 downregulates whereas KCC2 upregulates, leading to outward Cl^-^ directed transport. This allows a decline in the intracellular Cl^-^ concentration and a concomitant shift in GABA responses from depolarisation to hyperpolarisation (Ben-Ari, 2002; Ben-Ari *et al*., 2012). However, there are few adult neurones in the central nervous system that retain immature-like functional characteristics under physiological conditions with the absence of KCC2 expression and depolarising GABA transmission (Barthó *et al*., 2004; Haam *et al*., 2012; Sun *et al*., 2012)

Cerebrospinal fluid-contacting neurones (CSF-cNs) constitute a peculiar neuronal population present in all vertebrates spinal cord, from cyclostomes to mammals (Stoeckel *et al*., 2003*a*; Vígh *et al*., 2004; Orts-Del’immagine *et al*., 2012; Djenoune *et al*., 2014). They lie at the interface between CSF and the parenchyma while extending a dendrite towards the central canal (CC), ending with a terminal protrusion or ‘bud’ in contact with the fluid (Stoeckel *et al*., 2003*a*; Marichal *et al*., 2009; Orts-Del’immagine *et al*., 2012). Based on their morphology and localisation, it was proposed that CSF-cNs represent an intrinsic sensory module able to carry information from the CSF to spinal neuronal network (Fidelin *et al*., 2015; Böhm *et al*., 2016; Knafo *et al*., 2017; Gerstmann *et al*., 2022). Consistently, spinal CSF-cNs express selectively the polycystic kidney disease 2-like 1 (PKD2L1) channel, a sensory transduction protein sensitive to extracellular variation of pH, osmolarity and sensing CSF flow during spinal cord torsion (Orts-Del’immagine *et al*., 2012; Jalalvand *et al*., 2016; Orts-Del’Immagine *et al*., 2016; Sternberg *et al*., 2018). PKD2L1 are non-selective cationic channels and act as a potential generator in CSF-cNs for action potentials firing, suggesting a key role of PKD2L1 in tuning their excitability (Orts-Del’Immagine *et al*., 2016). CSF-cNs excitability is also adjusted by GABAergic synaptic inputs involving ionotropic GABA_A_-Rs as well as G protein-coupled metabotropic GABA_B_ receptors (GABA_B-_Rs) (Orts-Del’immagine *et al*., 2012; Jurčić *et al*., 2019). Despite these apparent hallmarks of functionally mature neurones, CSF-cNs were also shown to express molecular markers of immaturity even into adulthood, such as the migrating and neuronal shaping proteins doublecortin (DCX) and PSA‐NCAM (polysialylated neuronal cell adhesion molecule) as well as the homeobox protein Nkx6.1 (Stoeckel *et al*., 2003*b*; Shechter *et al*., 2007; Orts-Del’immagine *et al*., 2014). This suggests that CSF-cNs persist in a prolonged immature state in the adult spinal cord, though the functional relevance of such immaturity has not been addressed. Specifically, the KCC2/NKCC1 expression properties are unknown in CSF-cNs of adult animals and their GABA_A_ receptor-meditated transmission has, so far, only been studied with recording techniques that perturb the intracellular Cl^-^ concentration (Orts-Del’immagine *et al*., 2012; Jurčić *et al*., 2019)

In the present study, using immunostainings and confocal imaging, we show that CSF-cNs of adult mouse spinal cord express NKCC1 but not KCC2 transporters. Consistently, under Cl^-^ load with whole-cell patch-clamp recordings, CSF-cNs have a lower Cl^-^ extrusion capacity when compared to mature VGAT^+^ (GABA/glycine) neurones of spinal dorsal horn. Using non-invasive recording techniques for maintaining intact the internal Cl^-^ concentration, we further show that GABA_A_-Rs activation induces a predominant depolarising effect in adult spinal CSF-cNs. However, a fraction of CSF-cNs responds to GABA by a hyperpolarisation, suggesting distinct maturation states of CSF-cNs. Depolarising GABA_A_-Rs responses trigger action potentials in CSF-cNs but also shunting inhibition on their discharge activity. Calcium imaging experiments with the calcium sensor GCaMP6f selectively expressed in CSF-cNs revealed that GABA evokes intracellular calcium elevations dependent on the recruitment of voltage-gated calcium channels and intracellular chloride accumulation by NKCC1. Finally, as previously shown in medullar CSF-cNs (Jurčić *et al*., 2019), GABA_B_-Rs do not mediate G protein-coupled inwardly rectifying potassium (GIRK) currents in spinal CSF-cNs, suggesting that, in adult mice, CSF-cNs lack the hyperpolarising action of both GABA_A_-Rs and GABA_B_-Rs.

## Results

### Spinal CSF-cNs do not express KCC2 transporters in adult mice

CSF-cNs were shown to persist in an immature state in adult spinal cord (Orts-Del’immagine *et al*., 2014; Kútna *et al*., 2014). Moreover, it is well known that the neuron-specific chloride extruder KCC2 has a low activity in immature neurones of the CNS (Rivera *et al*., 1999; Blaesse *et al*., 2009). Therefore, we first investigated the expression of KCC2 in adult spinal CSF-cNs with immunostainings and using transgenic PKD-tdTomato mice expressing the fluorescent protein tdTomato in CSF-cNs for their identification. tdTomato-positive cells around the CC of lumbar spinal cord from adult mice had the typical morphology of CSF-cNs with small round soma (arrows, **Fig. 1A**) and short dendrites projecting towards the CC and ending with a bud (open arrowheads, **Fig. 1A**). Immunostainings against KCC2 revealed no detectable expression of the transporter in tdTomato-positive CSF-cNs (**Fig. 1B** and **C**; n = 3). In contrast, KCC2 immunoreactivity was observed in the plasmalemmal region of dorsal horn neurones as previously described in such mature neuronal population (arrowheads, **Fig. 1D**; n = 3) (Tang *et al*., 2015; Ferrini *et al*., 2020). Altogether, these data suggest a lack of KCC2 expression in spinal CSF-cNs of adult mice.

**Figure 1.**
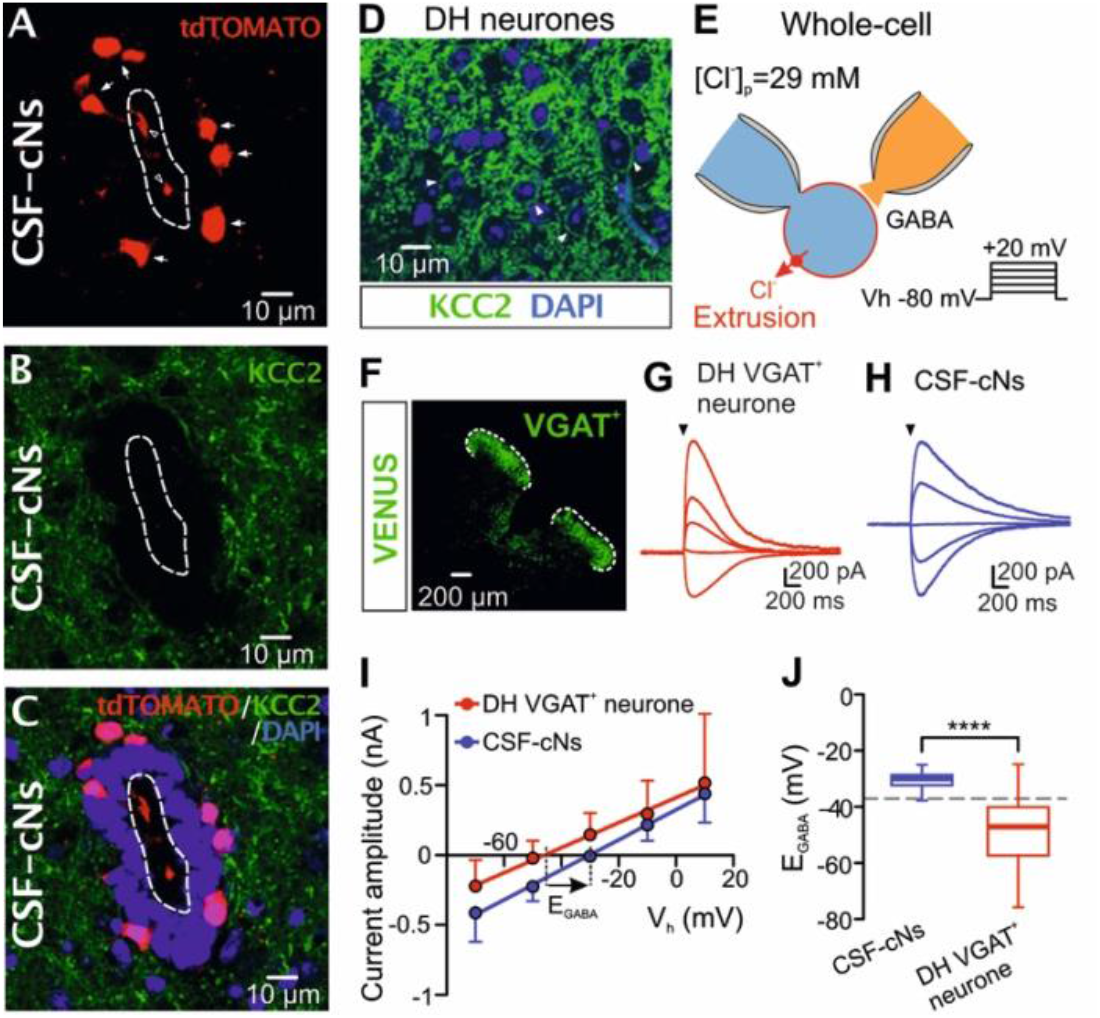
CSF-cNs of adult spinal cord do not express KCC2 and have low Cl-extrusion capacity. **A-C**, confocal microscopy images showing no detectable KCC2 immunostaining (green, panel B) in adult cerebrospinal fluid-contacting neurones (CSF-cNs) expressing the fluorescent protein tdTomato under the control of PKD2L1promoter (red, panel A; merged image in panel C). tdTomato fluorescence shows the typical morphology of CSF-cNs with a round soma (arrows) and a protrusion or bud ending their dendrites (open arrowheads). Cell nuclei were counterstained with DAPI (blue, panel C). The white dash line delineates the central canal (CC). **D**, confocal image showing plasmalemmal immunostaining for KCC2 (green, arrowheads) and DAPI nuclear staining (blue) in dorsal horn (DH) neurones of mouse spinal cord. **E**, Cl^-^ extrusion measurements were performed in whole-cell configuration by recording GABA_A_ receptors-mediated currents at different holding potentials (Vh: from -80 mV to +20 mV; steps of +10 mV) and with a high Cl_-_ pipette solution (29 mM). **F**, Confocal image showing VGAT^+^ (GABA/glycine) neurones of the spinal dorsal horn (white dash line) expressing the fluorescent Venus protein. **G-H**, representative averaged current traces evoked by puff applications (arrowheads) of GABA (1 mM, 30 ms duration) recorded in dorsal horn (DH) VGAT+ neurons (G, red traces) and CSF-cNs (H, blue traces) in the presence of DNQX (20 μM) and strychnine (1 μM). **I**, mean GABA I-V curves recorded from DH VGAT+ neurones (red) and CSF-cNs (blue). In CSF-cNs EGABA (−33.1 mV, R^2^ = 0.8; defined by the x-intercept of the linear fit, dashed line) is shifted to more positive values compared to DH VGAT^+^ neurones (−45.9 mV, R^2^ = 0.5). Membranepotentials were corrected for liquid junction potential (10 mV). **J**, box-and-whisker plot of E_GABA_ from CSF-cNs (−30.5 ± 3.4 mV, n = 18) and DH VGAT+ neurones (−47.8 ± 13 mV, n = 19). E_GABA_ in CSF-cNs was significantly more depolarised compared to DH neurones (****p = 1.6 × 10-5, unpaired Student’s t-test). The grey dashed line shows the theoretical E_GABA_ (−38 mV) calculated with GHK equation.

### Low Cl-extrusion capacity in spinal CSF-cNs

KCC2 deficiency may involve lower Cl^-^ extrusion capacity in CSF-cNs. To test this hypothesis, we compared the Cl^-^ extrusion capability between CSF-cNs and dorsal horn (DH) VGAT^+^ (GABA/glycine) neurones expected to have active KCC2-dependant Cl^-^ extrusion (Ferrini *et al*., 2020). Cl^-^ extrusion capacity was assessed under constant Cl^-^ load by recording CSF-cNs and DH VGAT^+^ neurones in whole-cell configuration with high Cl^-^ concentration in the recording pipette (29 mM, **Fig. 1E**). Under these conditions and according to the Goldman-Hodgkin-Katz (GHK), the theoretical value of the reversal potential of GABA_A_ receptor-mediated currents (E_GABA_) equation is -38 mV. GABA-mediated currents were elicited by pressure applications of GABA (1 mM, 30 ms) and recorded at different holding potentials to generate the IV curve and determine E_GABA_ (see Methods). The E_GABA_ determined experimentally was then compared to the calculated one. As previously described, a more negative imposed E_GABA_ compared to the calculated value reveals active Cl^-^ extrusion (Cordero-Erausquin *et al*., 2005; Schmidt *et al*., 2018; Ferrini *et al*., 2020). Using (VGAT)-venus transgenic mice allowing the expression of the venus protein in VGAT^+^ neurones (**Fig. 1F**), we recorded dorsal horn (DH) VGAT^+^ neurones and found they have an experimental value of E_GABA_ (−47.8 ± 13 mV, n = 19) hyperpolarised from the calculated E_GABA,_ indicating effective Cl^-^ extrusion (**Fig. 1G, I** and **J**). In CSF-cNs lacking KCC2, E_GABA_ was significantly more depolarised (−30.5 ± 3.4 mV, n = 18; Fig. 1H and I) compared to DH VGAT^+^ neurones (p = 1.6 × 10^−5^, unpaired Student’s t-test; **Fig. 1J**) and close to the calculated E_GABA_, hence indicating a weaker extrusion capacity in CSF-cNs. Consistently, bath application of the KCC2 inhibitor furosemide (100 μM) shifted E_GABA_ to more positive values in DH VGAT^+^ neurones (control: -45.6 ± 15.9 mV; FURO: - 27.6 ± 8.4 mV, n = 8; p = 0.0078, Wilcoxon matched-pairs signed rank test; **Fig. 2A, B** and **C**) but had no effects on E_GABA_ in CSF-cNs (control: -28.6 ± 4.1 mV; FURO: -27.7 ± 5.2 mV, n = 11; p = 0.5, Student’s two-tailed paired t-test; **Fig. 2D, E** and **F**). Collectively, these results show that KCC2 is a pivotal Cl^-^ extruder in mature DH GABA/glycine neurones whereas no KCC2 activity was observed in CSF-cNs.

**Figure 2.**
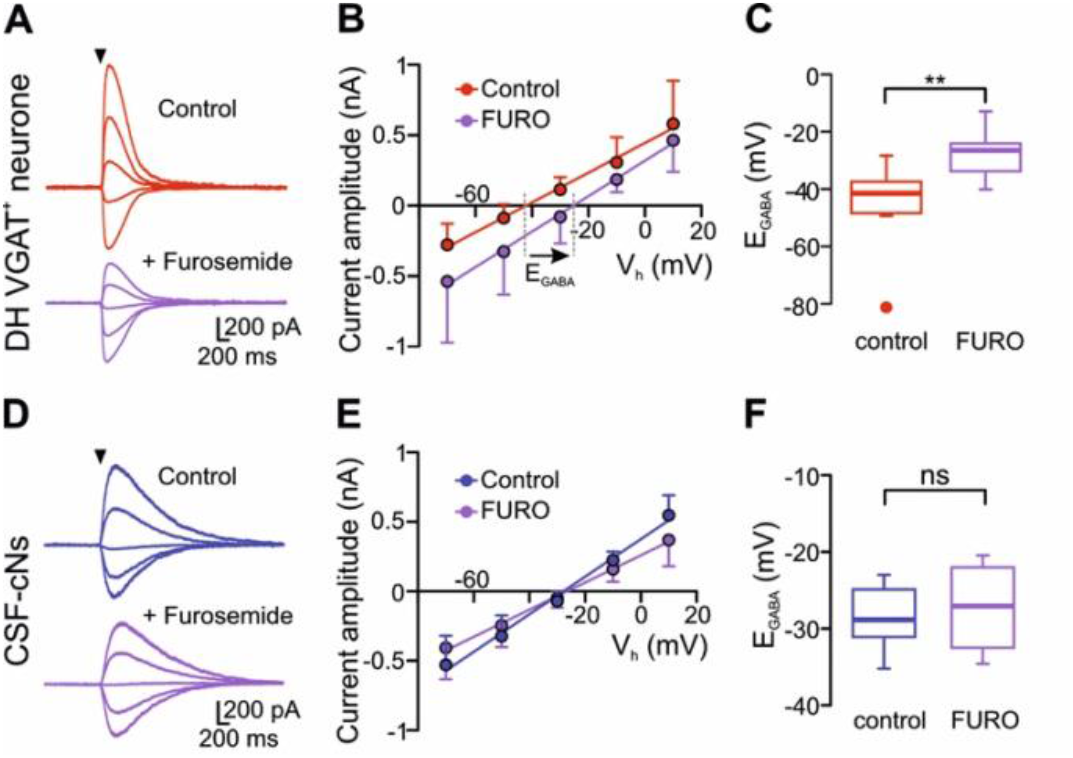
Inhibition of KCC2 shifts the E_GABA_ in DH VGAT^+^ neurones but not in CSF-cNs. **A and B**, representative current traces (A) and mean I-V plots (B) of GABA-induced currents recorded from DH VGAT^+^ neurones in control (red circles) and in the presence of the KCC2 inhibitor furosemide (FURO, 100 μM; purple circles). Currents were elicited with puff application of GABA (1mM for 30 ms; arrowheads) in the presence of DNQX (20 μM) and strychnine (1μM). E_GABA_ obtained with furosemide is shifted to more positive values compared to control (control: E_GABA_ = -42.1 mV, R^2^ = 0.7; FURO: E_GABA_ = -25,3 mV, R^2^ = 0.7; x-intercept of the linear fit, grey dashed line). **C**, summary box-and-whisker plot of E_GABA_ from DH VGAT^+^ neurones in control and furosemide conditions (control: -45.6 *±* 15.9 mV; FURO: -27.7 *±* 5.2 mV, n = 8; **p = 0.0078, Wilcoxon matched-pairs signed rank test). **D and E**, representative GABA-induced current traces (D) and mean I-V curves (E) from CSF-cNs in control (blue) and in the presence of 100 μM furosemide (purple). **F**, box-and-whisker plot showing that furosemide has no effect on E_GABA_ from CSF-cNs (control: -28.6 *±* 4.1 mV; FURO: -27.7 *±* 5.2 mV, n = 11; “ns”, no significant difference, p = 0.5, Student’s two-tailed paired t-test). Membrane potentials were corrected for liquid junction potential (10 mV).

### GABA_A_ receptors depolarise spinal CSF-cNs

Cl^-^ transmembrane extrusion by KCC2 results in hyperpolarising action of GABA_A_-Rs activation in mature neurones. Therefore, one could suggest that the absence of KCC2 activity in CSF-cNs contributes to accumulate intracellular Cl^-^, thus resulting in depolarising action of GABA_A_-Rs activation by Cl^-^ efflux through the receptor. To test this hypothesis, we investigated the effect of GABA_A_-Rs activation on the membrane potential of CSF-cNs recorded in the cell-attached configuration, thus preserving intact intracellular Cl^-^ concentration. We used the reversal potential of K^+^ channels (E_K_) as a read out for CSF-cNs resting membrane potential as previously described in other cell types (Verheugen *et al*., 1999; Fricker *et al*., 1999). K^+^ channels were activated with voltage-ramps from -140 to +200 mV (−V_pip_) at 1 s intervals and CSF-cNs membrane potential estimated by measuring E_K_ at the intersection between the K^+^ current and the fit to the linear component (**Fig. 3Aa**, red line). Among 23 CSF-cNs recorded, in the presence of DNQX (20 μM), strychnine (1 μM) and TTX (0.5 μM), pressure application of 1 mM GABA for 10 s induced a depolarisation of 19 ± 8.9 mV in 16 cells (control membrane potential: -61.1 ± 10.8 mV; membrane potential during GABA application: -42 ± 9.0 mV, n = 16; p = 3.7 × 10^−7^, Student’s two-tailed paired t-test; **Fig. 3Ba, Bb, D** and **F**). In the remaining 7 neurones, GABA induced a hyperpolarisation of 9 ± 5.5 mV (control: -44.4 ± 10.3 mV; with GABA: -53.4 ± 10.2 mV; n = 7; p = 0.0048, Student’s two-tailed paired t-test; **Fig. 3Ca, Cb, E** and **F**). Note that the control membrane potentials between the two groups of CSF-cNs (−61.1 ± 10.8 mV and -44.4 ± 10.3 mV for cells responding with depolarisation and hyperpolarisation, respectively) were significantly different (p = 0.0042, unpaired Student’s t-test). Finally, bath application of GABA_A_-Rs antagonists gabazine (10 μM) and picrotoxin (100 μM) blocked the GABA modulation of CSF-cNs membrane potential (p = 0.4, Student’s two-tailed paired t-test; **Fig. 3H**), thus demonstrating the involvement of GABA_A_-Rs activation in GABA effects.

**Figure 3.**
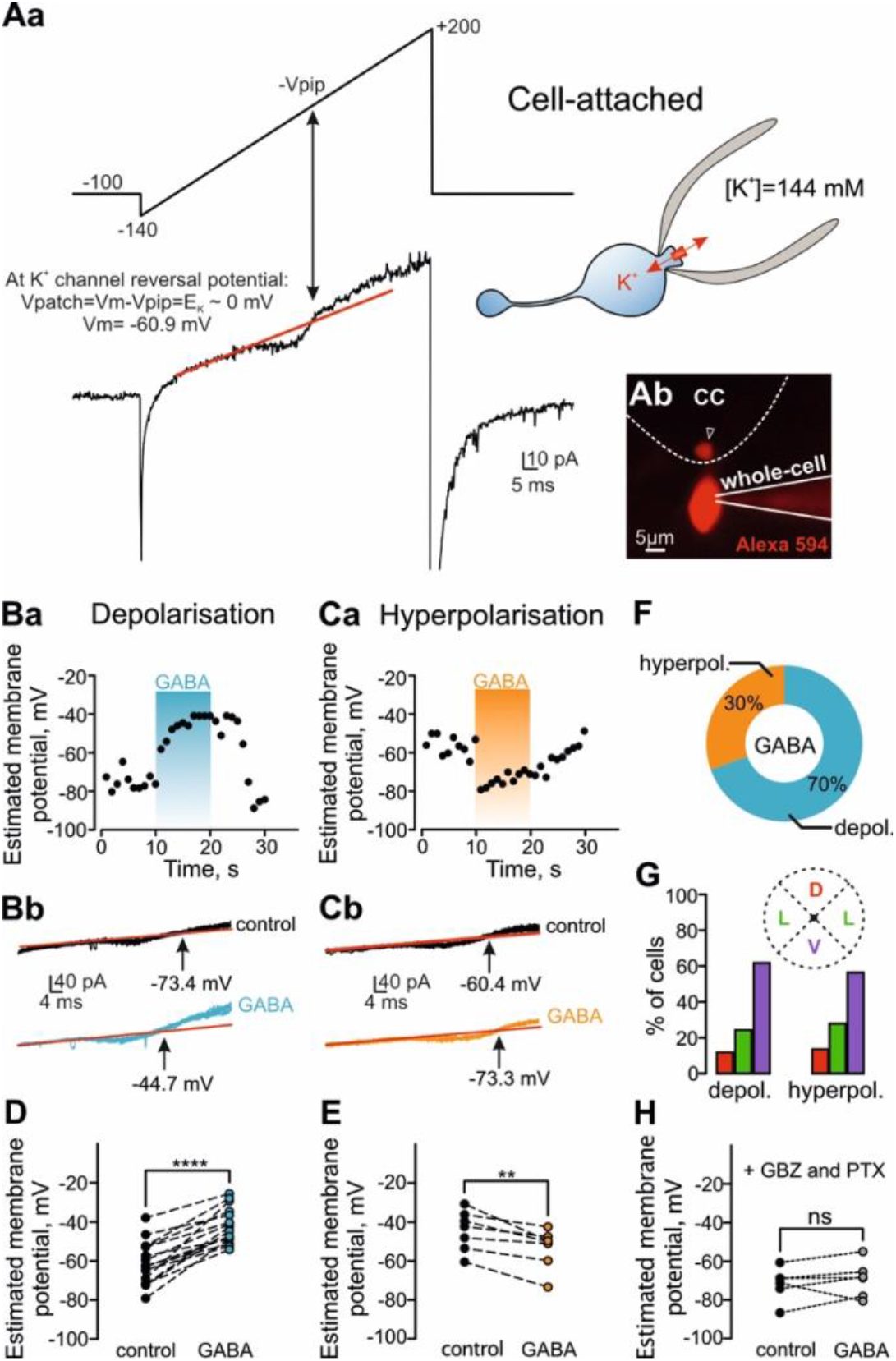
Activation of GABA_A_ receptors evokes a predominant depolarising effect in spinal CSF-cNs. **A**, estimation of membrane potential in intact CSF-cNs (a) was achieved by recording voltage-gated K^+^ currents with voltage-clamp tight-seal cell-attached configuration. Current (bottom trace) was evoked with a depolarising ramp command from -140 mV (−V_pip_) to +200 mV (top trace). The holding potential (−V_pip_) was -100 mV with respect to the cell membrane potential. K^+^ currents were recorded with 144 mM K^+^ pipette solution. The reversal of cell-attached K^+^ currents was determined from the intersection of the fit to the linear leak (red line) and the K^+^ current, then used to measure the cell membrane potential. CSF-cNs typical morphology was confirmed after whole-cell dialysis with Alexa 594 and the visualization of the bud (open arrowhead in b). The white dashed line delineates the central canal (CC). **B**, plot of the CSF-cNs membrane potential versus time (a) and 10 superimposed current traces (b) before (black traces) and during GABA application (blue traces) showing that GABA evoked a depolarisation. GABA was pressure applied (1mM, 10 s duration) and recording was performed in aCSF supplemented with DNQX (20 μM), strychnine (1 μM) and TTX (0.5 μM). CSF-cNs membrane potential in both conditions was measured as shown in (A). Note the shift of K^+^ current reversal (b) to the left with GABA (corresponding to a more depolarised membrane potential). **C**, time course of CSF-cNs membrane potential (a) and 10 superimposed current traces (b) in control (black) and during GABA application (orange) showing a hyperpolarising effect of the agonist. **D and E**, Line plots showing the membrane potential of individual CSF-cNs in control and during GABA application. GABA evoked in CSF-cNs a significant depolarisation (panel D; control: -61 ± 10.8 mV; GABA: -42 ± 9.0 mV, n = 16; ****p = 3.7 × 10^−7^, Student’s two-tailed paired t-test) and hyperpolarisation (panel E; control: -44.4 ± 10.3 mV; GABA:-53.4 ± 10.2 mV, n = 7; **p = 0.0048, Student’s two-tailed paired t-test). **F**, pie graph indicating the proportion of CSF-cNs responding to GABA with depolarisation (blue, 70%, n = 16) or hyperpolarisation (orange, 30%, n = 7). **G**, bar graph summarizing the distribution of CSF-cNs with depolarising or hyperpolarising responses along the dorsal-ventral axis of the CC. The region around the CC was subdivided in 4 quarters to determine the position of the recorded cell (inset, D: dorsal; L: lateral; V: ventral). **H**, line plot of individual CSF-cNs showing that GABA has no effect on their membrane potential in the presence of GABA_A_ receptors antagonists gabazine (GBZ, 10 μM) and picrotoxin (PTX, 100 μM) (control: -71.7 mV ± 8.6 mV; GABA: -69.3 ± 9.3 mV, n = 6; “ns”, no significant difference, p = 0.4, Student’s two-tailed paired t-test).

Altogether, these results revealed CSF-cNs as a heterogenous cell population in adult mice with different GABA_A_-Rs modulation supporting different state of maturation, the immature CSF-cNs subgroup with depolarising GABA signalling being the predominant population. Finally, as CSF-cNs with different maturation state were shown to have a different localisation around the CC (Petracca *et al*., 2016), we analysed the distribution of CSF-cNs that responded to GABA with either depolarisation or hyperpolarisation. As shown in **Fig. 3G**, there was no difference in the distribution of CSF-cNs with depolarising or hyperpolarising responses across dorsal, lateral, and ventral regions of the CC.

### Depolarising GABA_A_ receptors generate AP in CSF-cNs

We next examined the effect of depolarising GABA responses on CSF-cNs excitability. For each cell, we first determined the effect of GABA on the membrane potential (depolarisation or hyperpolarisation, **Fig 4Aa** and **Ba** or **Da**, respectively) with cell-attached recording in tight-seal current-clamp configuration. Subsequently in the same cell, we analysed in cell-attached voltage-clamp mode the GABA effects on CSF-cN action potential (AP) firing properties. (Perkins, 2006).

**Figure 4.**
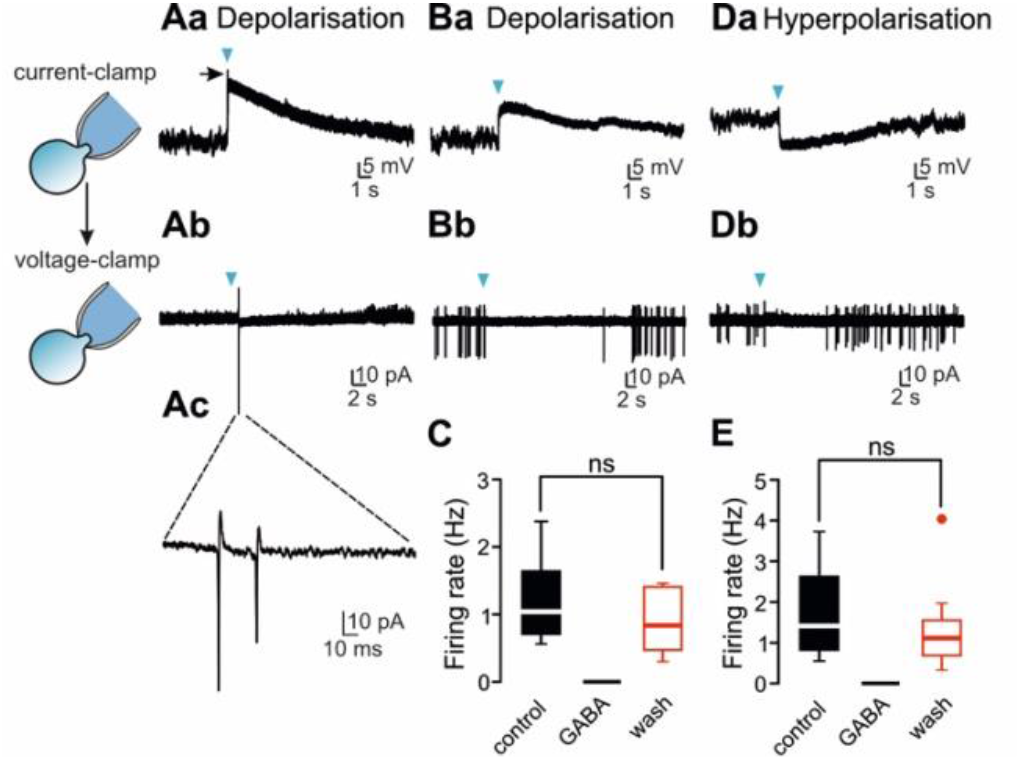
Depolarising GABA triggers AP or mediates shunting inhibition in CSF-cNs. **A**, pressure application of GABA (1 mM for 5 s, arrowheads) in the presence of DNQX (20 μM) and strychnine (1 μM) induced a depolarisation (from -47.4 mV to -29.1 mV; the arrow indicates the presence of an action potential) in CSF-cNs recorded in current-clamp cell-attached (a). In the same cell recorded in voltage-clamp cell-attached, GABA induced the discharge of action potential (AP) currents (b and c). **B**, CSF-cNs with GABA-mediated depolarisation (a, current-clamp cell-attached; from -47.6 mV to -36.6 mV) and an inhibition of the AP spontaneous activity following GABA application (b, voltage-clamp cell-attached). **C**, summary box-and-whisker plot showing that GABA abolished in a reversible manner the AP spontaneous activity of CSF-cNs that responded by depolarisation (control: 1.2 ± 0.6 Hz; wash: 0.9 ± 0.5 Hz, n = 6; “ns”, no significant difference, p = 0.1, Student’s two-tailed paired t-test). **D**, current-clamp (a) and voltage-clamp recordings (b) of a representative CSF-cNs that responded to GABA by a hyperpolarisation (a, from -44.7 mV to -54.2 mV) and a reduction of the firing frequency (b). **E**, summary box-and-whisker plot of AP spontaneous activity of CSF-cNs responding by hyperpolarisation in control, GABA and after washing out the agonist (control: 1.7 ± 1.2 Hz; wash: 1.3 ± 1.1 Hz, n = 10; “ns”, no significant difference, p = 0.3, Wilcoxon matched-pairs signed rank test).

In the presence of DNQX and strychnine, pressure application of GABA (1 mM for 5 s) induced a depolarisation of 12.1 ± 4.9 mV (from -49.8 ± 6.1 mV to -37.7 ± 10.3 mV) and led to the generation of AP (**Fig. 4Aa, Ab** and **Ac**; n = 6). However, we also observed a reversible inhibition of the AP spontaneous activity in CSF-cNs that responded to GABA with lower magnitude depolarisations (7.5 ± 5.1 mV; from -46.8 ± 3.0 to -39.3 ± 5.3 mV, n = 6; **Fig. 4Ba, Bb** and **C**). This shows that depolarising GABA differentially modulates CSF-cNs in adult spinal cord by supporting AP generation or shunting inhibition. As expected, in CSF-cNs responding to GABA with hyperpolarisations (9.1 ± 4.9 mV; from -46 ± 4.8 mV to -55.1 ± 8.7 mV; n = 10), the agonist induced a reversible inhibition of AP firing (**Fig. 4Da, Db** and **E**). Note that the resting membrane potentials measured in current-clamp cell-attached between cells responding with depolarisation or hyperpolarisation were similar (p = 0.3, unpaired Student’s t-test; see Discussion).

### GABA evokes cytosolic Ca^+2^ elevations dependent on voltage-gated Ca^2+^ channels and NKCC1 transporters

Previous studies have demonstrated that GABA_A_-Rs promote depolarisation-dependent elevations of intracellular calcium concentration ([Ca^2+^]_i_) in young neurones (Owens *et al*., 1996; Garaschuk *et al*., 2000; Reali *et al*., 2011). We thus investigated whether depolarising GABA_A_-Rs in spinal CSF-cNs of adult spinal cord induces changes in [Ca^2+^]_i_ using calcium imaging with the GCaMP6f sensor, selectively expressed in CSF-cNs by Cre/lox recombination (PKD-CCaMP6f mice, see Methods). In the presence of DNQX (20 μM), strychnine (1 μM) and CGP54626 (10 μM, a selective blocker of GABA_B_Rs), pressure application of GABA (1 mM, 10 s duration) induced an increase in [Ca^2+^]_i_ (79.5 ± 81.6 % ΔF/F_0_, n = 29) in CSF-cNs (**Fig. 5A, C** and **E**). By comparison, the depolarisation of CSF-cNs by application of aCSF solution with high K^+^ concentration (20 mM for 10 s) triggered similar [Ca^2+^]_i_ rise (93.4 ± 90.2% ΔF/F_0_, n = 23; p = 0.2, Mann Whitney-test; **Fig. 5B, D** and **E**). However, the proportion of GABA-responsive CSF-cNs was significantly lower compared to cells with KCl responses (GABA: 29/65, 45%; KCl: 23/26; 88%; p = 0.00013, Fisher’s exact test; **Fig. 5F**). Next, we studied the impact of different treatments on GABA-mediated [Ca^2+^]_i_ changes. As we experienced rapid bleaching of the dye preventing repetitive [Ca^2+^]_i_ measurements in the same cell, we determined the proportions of CSF-cNs that respond to GABA by [Ca^2+^]_i_ rise under different experimental conditions and compared it to control (GABA alone). As shown in **Fig. 5F**, in the presence of aCSF solution lacking Ca^2+^ (0-Ca^+2^) or containing the voltage-dependent calcium channel blocker cadmium (Cd^+2^, 200 μM), the proportion of CSF-cNs showing [Ca^2+^]_i_ increase following GABA application was significantly lower (0-Ca^+2^: 2/20, 10%, p = 0.007; Cd^2+^: 7/44, 15%, p = 0.002; Fisher’s exact test). Moreover, the inhibition of the intracellular Cl^-^ accumulator NKCC1 by the selective blocker bumetanide (BTN, 20 μM) significantly reduced the proportion of responding CSF-cNs compared to control (7/37, 19%; p = 0.010, Fisher’s exact test, **Fig. 5F**). In line with this finding, immunohistochemistry analysis revealed NKCC1 staining in spinal CSF-cNs (open arrowheads, **Fig. 5G, H** and **I**). Altogether, these results show that NKCC1 accumulates intracellular Cl^-^ in CSF-cNs allowing depolarising GABA_A_ receptors-mediated calcium influx through voltage-gated calcium channels.

**Figure 5.**
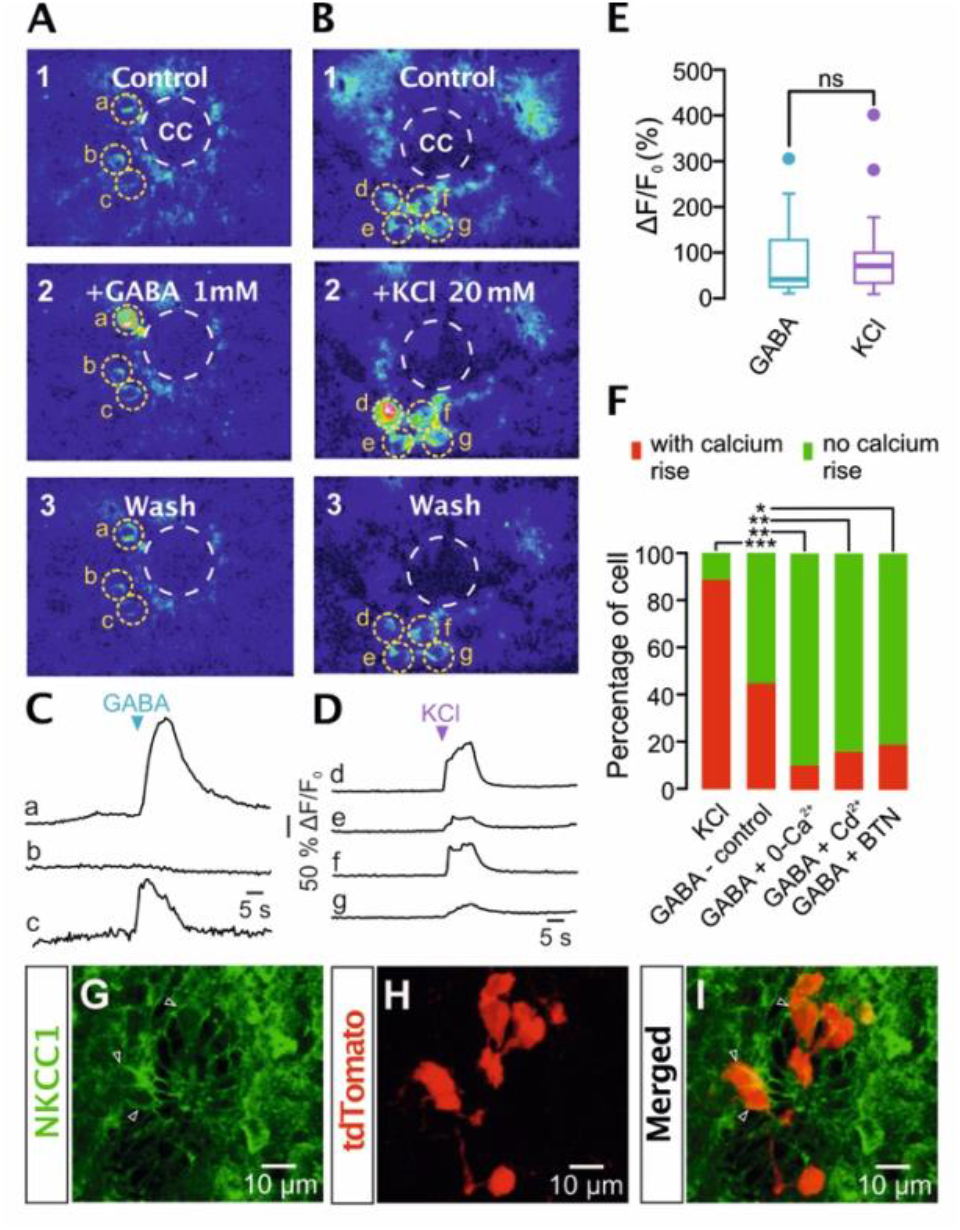
Activation of GABA_A_ receptors evokes calcium transients in CSF-cNs. **A and B**, representative images illustrating the baseline fluorescence (1), the maximum fluorescence increase after pressure application of 1 mM GABA or 20 mM KCl (2) for 10 s and after the recovery period (3, wash) in CSF-cNs expressing the calcium sensor GCaMP6f. GABA and KCl were applied in aCSF supplemented with DNQX (20 μM), strychnine (1 μM) and CGP54626 (10 μM). Images are presented in pseudocolour (black: low fluorescence; red: high fluorescence). Yellow dashed lines indicate the regions of interest (ROI) drawn on CSF-cNs soma. Dashed white lines delineate the central canal (CC). **C and D**, time course of the net fluorescence changes during application of GABA (C) and KCl (D) from cells shown in (A) and (B), respectively. Note that GABA failed to induce a fluorescence rise in cell (b). **E**, box-and-whisker plot for the normalized fluorescence variation (ΔF/F_0_) induced by GABA and KCl in CSF-cNs (GABA: 79.5 ± 81.6 %, n = 29; KCl: 93.4 ± 90.2%, n = 23; “ns”, no significant difference, p = 0.2, Mann Whitney-test). **F**, stacked bar-graph showing the percentage of CSF-Ns that responded by an increase in [Ca^2+^]_i_ to KCl and GABA under different perfusion conditions: aCSF without calcium (0-Ca^2+^) and aCSF supplemented with the voltage-gated Ca^2+^ channel blocker cadmium (Cd^2+,^ 200 μM) or bumetanide (BTN, 20 μM), a selective inhibitor of NKCC1 transporter (***p = 0.00013, KCl compared to control GABA; **p = 0.007, ** p = 0.002, *p = 0.010, GABA control compared respectively to 0-Ca^2+^, Cd^2+^ and BTN, Fisher’s exact test). **G-I**, confocal fluorescence images of histological sections showing NKCC1 immunostaining (green, open arrowheads, panel G) in tdTomato-expressing CSF-cNs (red, panel H; merged image in panel I).

### No hyperpolarising modulation by GABA_B_ receptors in spinal CSF-cNs

Medullo-spinal CSF-cNs in adult rodents were shown to express metabotropic G protein-coupled GABA_B_-Rs (Margeta-Mitrovic *et al*., 1999; Jurčić *et al*., 2019). As hyperpolarising signalling of GABA_B_-Rs depends on the gating of inwardly rectifying K^+^ (GIRK) channels, we thus tested whether activation of GABA_B_-Rs generates outward K^+^ currents in spinal CSF-cNs that exhibit depolarising GABA_A_-Rs responses. The experimental design consisted of recording CSF-cNs first in current-clamp cell-attached configuration to determine the effect of GABA on the membrane potential without perturbation of intracellular Cl^-^ and subsequently, we established a whole-cell configuration of the same cell and recorded in voltage-clamp mode at a holding potential of -50 mV, the effect of GABA_B_-Rs activation on the holding current. We found that in CSF-cNs with depolarising GABA responses recorded in cell-attached (**Fig. 6Aa**), bath application of the selective GABA_B_-Rs agonist baclofen (100 μM) had no effects on the holding current (**Fig. 6Ab**; n = 6). These results indicate that CSF-cNs with depolarising GABA_A_-Rs do not express GABA_B_Rs-dependent K^+^ currents and therefore show CSF-cNs as a unique spinal neuronal population with neither GABA_A_- Rs nor GABA_B_-Rs hyperpolarising signalling.

**Figure 6.**
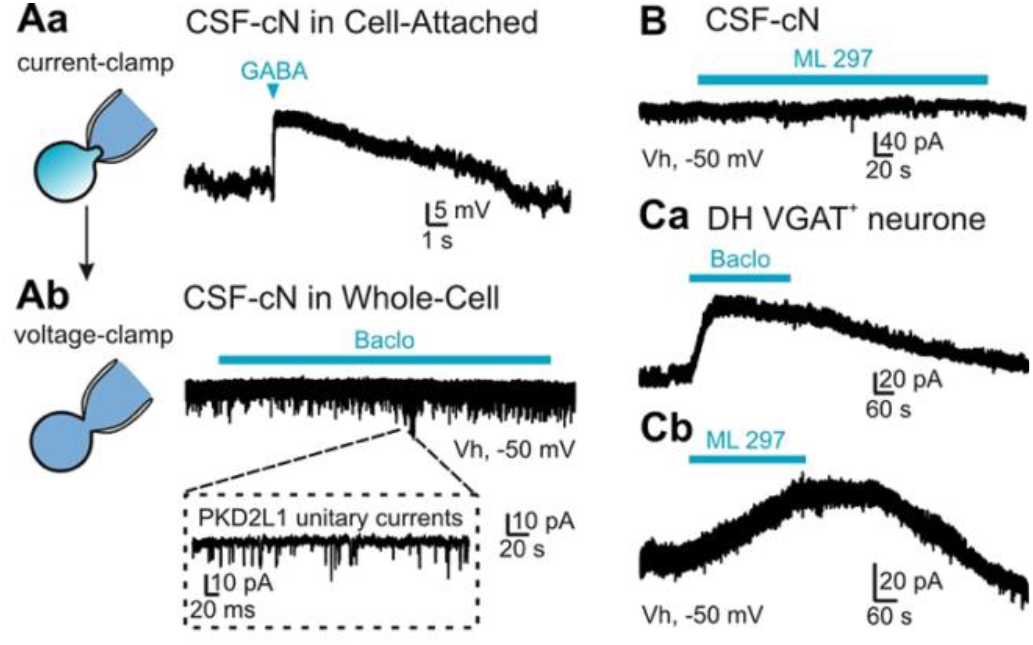
Spinal CSF-cNs do not express GABA_B_ receptors-mediated (GIRK) potassium currents. **A**, CSF-cNs recorded in current-clamp cell-attached (a) showing a depolarisation of the membrane potential following puff-applied GABA (1mM, 5 s). In the same cell illustrated in (a) voltage-clamped at -50 mV in whole-cell configuration, bath application of the selective agonist of GABA_B_ receptors baclofen (100 μM) had no effect on the holding current (b). Note the single-channel activity (inset) mediated by the PKD2L1 channel, as previously described in CSF-cNs. **B**, bath application of the GIRK1/2 activator ML297 (100 μM) did not generate K^+^ currents in CSF-cNs recorded with voltage-clamp whole-cell at a holding potential of -50 mV. **C**, in dorsal horn (DH) VGAT^+^ neurones recorded with voltage-clamp whole-cell at a holding potential of -50 mV, both baclofen (a) and ML297 (b) induced an outward K^+^ current. Baclofen and ML297 were bath applied at a concentration of 100 μM.

We further investigated the mechanisms underlying the lack of GABA_B_ receptors-dependent K^+^ currents, assuming that GIRK channels are not expressed in spinal CSF-cNs. To test this hypothesis, we applied on spinal CSF-cNs ML297, a direct and selective activator of GIRK1/2 channels, the most abundant neuronal GIRK channels (Gonzalez *et al*., 2018; Kimura *et al*., 2020; Luo *et al*., 2022). However, ML297 failed to elicit an increase in the holding currents in CSF-cNs (**Fig. 6B**; n = 11). In contrast and in agreement with previous reports (Kangrga *et al*., 1991; Kimura *et al*., 2020), bath application of baclofen (100 μM) or ML297 (100 μM) induced an outward K^+^ currents in DH VGAT^+^ neurones with amplitudes of respectively 38.8 ± 22.2 pA (n = 12) and 34.8 ± 21.2 pA (n = 7) (**Fig. 6Ca** and **Cb**), which confirms the potent modulator effect of ML297 on GIRK channels. Therefore, the deficit in GIRK expression probably supports the lack of GABA_B_ ^-^Rs-mediated hyperpolarisation in spinal CSF-cNs.

## Discussion

Immature neurones in adult mammalian central nervous system (CNS) represent a potential reservoir of young cells that may be mobilized under physiological or pathological conditions, such as injury or disease (La Rosa *et al*., 2019). Immature neurones were generally thought to localize in the subgranular zone of the hippocampal dentate gyrus and the subventricular zone of the lateral ventricles, but recent studies have suggested the presence of immature neurones on a large scale in other parts of the CNS as well, such as the spinal cord and the hindbrain (Rusanescu & Mao, 2014; Rusanescu, 2016). Specifically, an intriguing expression of immature neuronal markers has been shown in medullo-spinal CSF-cNs of adult rodents (Stoeckel *et al*., 2003*b*; Orts-Del’immagine *et al*., 2014; Kútna *et al*., 2014), but the functional significance of such immaturity was not yet known. In the present study, we demonstrate that GABA, acting via GABA_A_-Rs, predominantly depolarises spinal CSF-cNs, which triggers AP or shunts their tonic activity. In agreement with these results, we found that CSF-cNs express the NKCC1 transporter but lack KCC2, suggesting NKCC1-mediated intracellular Cl^-^ accumulation in CSF-cNs. Our study further shows that GABA recruits voltage-gated calcium channels to increase [Ca^2+^]_i_ in CSF-cNs. Finally, we found that CSF-cNs don’t express G protein-activated inwardly rectifying K^+^ (GIRK) currents, resulting in the absence of GABA_B_ receptor-mediated inhibition.

### Immature GABA_A_-Rs signalling in CSF-cNs of adult spinal cord

Previous studies with intracellular recording techniques have shown that CSF-cNs in adult mice receive GABAergic synaptic inputs that have been thought to mediate GABA_A_ receptor-dependent inhibition (Orts-Del’immagine *et al*., 2012; Jurčić *et al*., 2019). By using non-invasive cell-attached recordings for maintaining intact the intracellular milieu, we show instead that GABA predominantly depolarises CSF-cNs of adult spinal cord by acting on GABA_A_-Rs. A similar observation was previously made in spinal CSF-cNs of postnatal rats and juvenile turtle (Marichal *et al*., 2009; Reali *et al*., 2011), though our data are the first showing recordings of depolarising GABA responses in fully adult animals. Depolarising GABA signalling in CSF-cNs may imply a differential expression profile between the Cl^-^ extruder KCC2 and the Cl^-^ loader NKCC1 as described in immature neurones (Kilb, 2021). Indeed, immunohistochemical and functional analysis indicate a lack of KCC2 expression and a consistent low capacity of CSF-cNs for extruding Cl^-^ by imposing a constant Cl^-^ load through the recording pipette. In contrast, we observed in neurones of the spinal dorsal horn, considered as mature cells, a clear KCC2 immunostaining as well as an active Cl^-^ extrusion mediated by KCC2 transport under Cl^-^ load, as previously described (Cordero-Erausquin *et al*., 2005; Ferrini *et al*., 2020). In addition to the apparent low capacity of CSF-cNs to extrude Cl^-^, we found with immunohistofluorescence that CSF-cNs express the Cl^-^ accumulator NKCC1. Therefore, one could suggest that the depolarising GABA responses recorded in CSF-cNs are the result of a differential expression of Cl^-^ transporters, NKCC1 being predominant over KCC2, thus allowing high intracellular Cl^-^ concentrations in CSF-cNs (Rivera *et al*., 1999; Kilb, 2021). Consistent with this interpretation, blockade of NKCC1 with bumetanide reduced the proportion of CSF-cNs responding with intracellular calcium elevation evoked by GABA-dependent depolarisations. This latter observation also supports the point that GABA_A_ receptor-dependent depolarisations primarily result from a depolarising Cl^-^ flux rather than bicarbonate (Kaila *et al*., 1993). Additionally, the differential expression of NKCC1 and KCC2 in CSF-cNs from perfused spinal cord suggests that our electrophysiological data showing GABA_A_-Rs depolarising responses are not the result of a selective change in Cl^-^ regulation by the slicing procedure.

### Different state of maturation for spinal CSF-cNs in adult mice?

In addition to a primarily depolarising effect of GABA_A_-Rs, we observed GABA-mediated hyperpolarisations in CSF-cNs with cell-attached recordings, suggesting that CSF-cNs in the adult spinal cord are present around the CC in two putative distinct maturational states, immaturity with depolarising GABA_A_-Rs responses and a more mature state with hyperpolarising modulation. This observation is supported by previous studies showing that subgroups of CSF-cNs have different expression levels of the neuronal nuclear protein (NeuN), a marker of neuronal differentiation and maturation (Orts-Del’immagine *et al*., 2014; Orts-Del’Immagine *et al*., 2017). Moreover, CSF-cNs were shown to originate from distinct progenitors and distribute differently around the CC, with immature neurones located ventrally to the CC adopting single spike activity, whereas more mature neurones in the lateral position have repetitive spiking (Marichal *et al*., 2009; Petracca *et al*., 2016). This contrasts with our findings showing no segregated CSF-cNs around the CC with either depolarising or hyperpolarising responses, but rather a similar distribution of cells with the two types of GABA responses along the dorsal-ventral axis of the CC. It should be noted however that the differential localisation around the CC between immature and mature CSF-cNs was observed in postnatal mouse spinal cord (Petracca *et al*., 2016), whereas our study was performed in adult animals. Thus, the neuronal reorganisation around the CC in the developing spinal cord might explain opposite results (Kútna *et al*., 2014).

The fact that we identified CSF-cNs with hyperpolarising GABA is surprising insofar as KCC2 staining was not detected in CSF-cNs. One possible explanation may be that other types of K^+^-Cl^-^ cotransporters participate in reducing the intracellular Cl^-^ concentration in CSF-cNs thus allowing the switch between depolarising GABA and hyperpolarisation modulation. For instance, KCC3 is expressed in the CNS, functions under isotonic conditions as KCC2 and was shown to promote Cl^-^ extrusion in adult sensory neurones (Race *et al*., 1999; Lucas *et al*., 2012). Further investigations using immunostainings or transcripts analysis will address the possible expression of KCC3 or other K^+^-Cl^-^ extruders in CSF-cNs.

The presence of two subgroups of CSF-cNs is also supported by the observation that CSF-cNs with opposite GABA responses have different resting membrane potentials. Notably, by measuring the reversal potential of K^+^ currents in voltage-clamp cell-attached, we estimated the resting membrane potential of CSF-cNs and showed that neurones with depolarising GABA responses have a significantly more negative membrane potential compared to cells with hyperpolarising GABA. In contrast, the membrane potential measured with current-clamp cell-attached in the two groups of CSF-cNs (with depolarising or hyperpolarising GABA responses) have similar values. However, the accuracy of the voltage measured in current-clamp cell-attached depends upon the values of resistors (Rseal, Rpatch and Rcell) making up the electrical circuit involved in the current-clamp configuration (Perkins, 2006). Therefore, the measurement preformed in voltage-clamp cell-attached with the reversal potential of K^+^ current is a more accurate method for estimating the membrane potential as the voltage measured in current-clamp will be always an underestimate of the actual magnitude of the resting membrane potential (Perkins, 2006). The mechanisms underlying the difference in the resting membrane potential between CSF-cNs with depolarising or hyperpolarising GABA remain unknown but one could suggest that the apparent maturation of CSF-cNs is accompanied by changes in the membrane expression of ionic channels (Gao & Ziskind-Conhaim, 1998; Carleton *et al*., 2003). Finally, it should be noted that our data showing a more negative membrane potential in CSF-cNs with GABA depolarising responses contrast with previous results obtained from intracellular recordings and correlating immaturity of neurones with a strongly depolarised resting membrane potential compared to adult neurones (Ben-Ari *et al*., 1989; LoTurco *et al*., 1991; Zhang *et al*., 1991; Spigelman *et al*., 1992; Psarropoulou & Descombes, 1999; Luhmann *et al*., 2000). However, membrane potentials measured in these studies using whole-cell patch-clamp or perforated-patch have been probably underestimated notably because of a seal conductance at the membrane-pipette interface that depolarises the membrane potential (Chavas & Marty, 2003; Tyzio *et al*., 2003). Consistently, membrane potentials in hippocampal immature neurones appeared to be more negative than initially thought when determined with a similar method to that performed in our study but using the reversal potential of N-methyl-d-aspartate (NMDA) channels in cell-attached recordings (Tyzio *et al*., 2003).

### Dual excitatory/inhibitory action of GABA_A_-Rs on CSF-cNs

The main question behind the finding of a depolarising GABA signalling is whether it generates an increase in neuronal excitability (Kilb, 2021). With voltage-clamp cell-attached configuration allowing recordings of AP activity (Perkins, 2006), we found that GABA mediates an excitatory effect in CSF-cNs by triggering AP or an inhibition of their spontaneous firing activity. It was proposed that the relation between E_GABA_ and AP threshold determines whether activation of GABA_A_-Rs mediates excitation (Ben-Ari, 2002; Kilb, 2021; Lombardi *et al*., 2021). Such considerations may explain the distinct effect of GABA observed with current-clamp cell-attached recordings in CSF-cNs expressing similar resting potentials (Fig. 4A and C), that is either AP are triggered by depolarising GABA (E_GABA_ more positive to AP threshold) or they aren’t (E_GABA_ more negative to AP threshold). Furthermore, we observed that in the cell where a subthreshold depolarisation was evoked, GABA application abolished the spontaneous AP activity (Fig. 4C and D). This inhibitory effect of depolarising GABA may reflect a shunt inhibition, the depolarising current mediating a decrease in the membrane resistivity thus rendering the cell less responsive to other excitatory inputs (Ben-Ari, 2002; Kilb, 2021). Alternatively, depolarising GABA can also inactivate sodium channels needed for action potential generation, thus promoting transient inhibition through this mechanism (Ben-Ari, 2002). As glutamatergic receptors were blocked in our study and synaptic activity operating at low rate in CSF-cNs (Orts-Del’immagine *et al*., 2012), the spontaneous discharge of AP observed in cell-attached records is probably mediated by PKD2L1, the large unitary conductance of the channel being sufficient to generate spikes (Orts-Del’Immagine *et al*., 2016). Therefore, one could suggest an interesting mechanism involving depolarisation mediated by GABA_A_-Rs for the regulation of the excitation provided by PKD2L1.

Determining E_GABA_ value in CSF-cNs is therefore of primary importance for better understanding the conditions of an excitatory or inhibitory GABA. E_GABA_ can be classically assessed by using gramicidin-perforated patch recordings (Akaike, 1996) but it was not possible to accomplish such experiments in our preparation. The fact that we conducted our experiments with spinal cord slices from adult animals may have restricted the potential of gramicidin to form prototypical ion channels as this membrane-perforating agent was used mostly in neuronal preparations from postnatal or young animals. Another method to determine E_GABA_ in intact cells consists to measure E_GABA_ using single NMDA and GABA_A_-receptor channel recordings in cell-attached configuration (Tyzio *et al*., 2003; Quilichini *et al*., 2012). However, CSF-cNs were shown to lack NMDA receptors (Orts-Del’Immagine *et al*., 2016; Jurčić *et al*., 2019) and the determination of E_GABA_ requires a sequential recording of unitary NMDA and then GABA_A_ currents by re-patching the cell, hence together rendering this approach unrealisable in CSF-cNs. Alternatively, one could consider an indirect determination of E_GABA_ by measuring the intracellular concentration of Cl^-^ in CSF-cNs using Cl^-^ sensitive fluorescent indicators (Kovalchuk & Garaschuk, 2012).

### GABA_A_-Rs and Ca^2+^: a tandem involved in the maturation of CSF-cNs?

GABA operates as a signal of maturation in a wide range of developing brain structures and its action was shown to depend upon an increase in [Ca^2+^]_i_ concentration induced by membrane depolarisation (Leinekugel *et al*., 1995; Eilers *et al*., 2001; Ben-Ari, 2002; Ge *et al*., 2006). By imaging intracellular Ca^2+^ in CSF-cNs using the Ca^2+^ sensor GCaMP6f, we discovered that a large fraction of CSF-cNs responded to GABA application by increasing [Ca^2+^]_i_. The percentage of CSF-cNs responding with [Ca^2+^]_i_ rise was significantly reduced when CSF-cNs were perfused with an external solution lacking Ca^2+^ or supplemented with cadmium, thus suggesting that GABA-dependent [Ca^2+^]_i_ increases required extracellular Ca^2+^ and activation of voltage-gated Ca^2+^ channels. However, as a proportion of CSF-cNs still showed a GABA-induced [Ca^2+^]_i_ elevation in the presence of cadmium, we cannot exclude alternative mechanisms underlying the GABA-dependent [Ca^2+^]_i_ increase in CSF-cNs such as osmotic tension created by GABA_A_-Rs activation (Chavas *et al*., 2004). Medullo-spinal CSF-cNs were shown to express both high-(N-type) and low-voltage (T-type) Ca^2+^ channels (Marichal *et al*., 2009; Jurčić *et al*., 2019; Johnson *et al*., 2022). Thus, even without generating action potentials, depolarising GABA may still induce [Ca^2+^]_i_ rise in CSF-cNs through the activation of T-Type Ca^2+^ channels, as observed in other neuronal types (Fukuda *et al*., 1998; Sun *et al*., 2012). As voltage-gated Ca^2+^ channels play a key role in neuronal differentiation (Desarmenien & Spitzer, 1991; LoTurco *et al*., 1995; Rusanescu *et al*., 1995), it is tempting to speculate that GABA_A_-Rs couple with Ca^2+^ for driving maturation and integration of CNF-cNs into the spinal circuit. Further research will clarify the detailed mechanisms and functional relevance of GABA-induced [Ca^2+^]_i_ elevations in spinal CSF-cNs.

### The absence of GABA_B_ receptors-mediated GIRK currents as additional evidence of CSF-cNs immaturity

GABA inhibition in the CNS is also mediated by the activation of GABA_B_-Rs, which recruits G-protein-activated inwardly rectifying K^+^ (GIRK) channels and generates hyperpolarisation (Pinard *et al*., 2010). However, in CSF-cNs expressing depolarising GABA_A_-Rs responses, we additionally showed that the selective agonist of GABA_B_-Rs baclofen does not activate GIRK currents. Remarkably, this indicates that CSF-cNs form a unique neuronal population in adult spinal cord with neither GABA_A_-Rs nor GABA_B_-Rs hyperpolarising signalling. As a control, we tested the effect of baclofen on VGAT^+^ dorsal horn neurones and found that the agonist induced classical outward K^+^ currents thus ruling out eventual bias in our recording’s conditions. Further, we found that ML297, the selective activator of heteromeric GIRK 1/2 channel (the most abundant GIRK channels in the CNS), induced K^+^ currents in VGAT^+^ dorsal horn neurones but not in CSF-cNs. As functional GABA_B_-Rs are present in CSF-cNs (Margeta-Mitrovic *et al*., 1999; Jurčić *et al*., 2019), one could suggest that the absence of GIRK currents in spinal CSF-cNs results from the lack of GIRK channels expression. However, even though heterotetramers GIRK channels containing GIRK1 and GIRK2 subunits predominate in the CNS, we cannot exclude the expression in CSF-cNs of GIRK channels assembled with other subunits such as GIRK3 or GIRK4 (Luo *et al*., 2022). As pharmacological agents that selectively activate GIRK3 or GIRK4-containing channels are missing, dedicated experiments using immunostainings or transcriptomic analysis will address this eventuality.

GABA_B_-Rs are expressed and functional early in development, though they are not yet involved in regulating neuronal excitability by gating GIRK channels in most developing brain regions (Gaiarsa & Porcher, 2013). Indeed, GIRK inhibition occurs in neurones after several weeks of postnatal maturation, GIRK channels being expressed and functionally coupled with GABA_B_-Rs the latest between the third and the fifth postnatal week in the brain (Nurse & Lacaille, 1999; Lei & McBain, 2003). As we conducted our experiments with 7-10 weeks old mice, thus beyond the expected maturation period of GIRK channels, the absence of GABA_B_ receptor-mediated GIRK inhibition further supports the notion that CSF-cNs are under development in adult mice. Interestingly, in developing brain regions where GABA_B_-Rs do not trigger GIRK activation, the receptor was shown to engage intracellular signalling pathways involved in neuronal development (Bony *et al*., 2013; Gaiarsa & Porcher, 2013; Giachino *et al*., 2014). Further investigations might reveal the possible recruitment of such signalling pathways by GABA_B_-Rs in CSF-cNs and whether it contributes to their maturation.

## Conclusion

In the present study, functional analysis using technical approaches preserving CSF-cNs integrity shows that GABA_A_-Rs mediate depolarisations in most of spinal CSF-cNs of adult mice as well as calcium elevations through the activation of voltage-gated Ca^2+^channels. Additionally, GABA_B_-Rs do not mediate inhibition because of an apparent deficit in GIRK channel expression in spinal CSF-cNs. Hence, we present here the first evidence that CSF-cNs of adult mice have GABAergic signalling hallmarks that support their immaturity state. Our results are in line with previous assumptions suggesting CSF-cNs are immature neurones in adult mice as they express markers being involved in cell migration and neurite outgrowth such as PSA-NCAM and DCX (Stoeckel *et al*., 2003*b*; Shechter *et al*., 2007; Orts-Del’immagine *et al*., 2014). On the other hand, CSF-cNs were shown to take part in advanced physiological functions as they cooperate with other spinal neurones for the regulation of motor activity and posture in mice (Gerstmann *et al*., 2022). This suggests that immature CSF-cNs terminate their development and integrate into existing spinal circuits, though the underlying mechanisms and how GABA signalling may tune such functional maturation of CSF-cNs remain to be elucidated. Another remaining question is the source of GABA that depolarises CSF-cNs. An interesting recent finding indicates that CSF-cNs are interconnected (Gerstmann *et al*., 2022), CSF-cNs regulating each other’s through reciprocal GABAergic connections, as they are themselves GABAergic neurones (Shechter et *al*., 2007; Orts-Del’immagine *et al*., 2014). Therefore, one will address whether activation of CSF-cNs generates depolarisations in synaptically connected CSF-cNs partners. Additionally, GABA is known as a neuromodulator circulating in the CSF and therefore, a nonsynaptic GABA modulation of CSF-cNs cannot be excluded (Liu *et al*., 2005; Bjorefeldt *et al*., 2018).

## Materials and methods

### Ethical approval

All experiments were conducted in conformity with the rules set by the EC Council Directive (2010/63/UE) and the French “Direction Départementale de la Protection des Populations des Bouches-du-Rhône (DDPP13)” (Project License Nr: APAFIS 17596; 2018111919329153. N.W. and License for the Use of Transgenic Animal Models Nr: DUO-5214). Protocols used agree with the rules set by the Comité d’Ethique de Marseille, our local Committee for Animal Care and Research. All animals were housed at constant temperature (21°C), in an enriched environment, under a standard 12h light-12h dark cycle, with food (pellet AO4, UAR, Villemoisson-sur-Orge, France) and water provided *ad libitum*. Every precaution was taken to reduce to the minimal the number of animals used and minimize animal stress during housing and prior to experiments.

### Animals

Wildtype (C57BL/6J) and transgenic adult mice (7-10 weeks old) of both sexes were used in this study. Electrophysiological experiments were performed using wildtype C57BL/6J and (VGAT)-venus transgenic mice that express the fluorescent protein Venus under the control of the mouse VGAT (vesicular GABA transporter) promoter (generous gift of Dr Y. Yanagawa, Okazaki, Japan) (Wang *et al*., 2009). This allowed the identification of GABA/glycine (VGAT^+^) mature neurones located in the spinal dorsal horn (see Results). For immunostainings experiments, PKD-tdTomato mice, used to identify spinal CSF-cNs, were generated by crossing PKD2L1-IRES-Cre transgenic mice (generous gift from Dr E. R. Liman) with transgenic reporter Rosa26-loxP-stop-loxP-tdTomato mice (line Ai14 from The Jackson Laboratory). Calcium imaging experiments were carried out in PKD-CCaMP6f mice obtained by breeding PKD2L1-IRES-Cre with transgenic reporter Rosa-GCaMP6f animals (line Ai95D supplied by The Jackson Laboratory), thus allowing the selective expression of the GCaMP6 fast variant calcium sensor in CSF-cNs (Madisen *et al*., 2015).

### Immunohistochemistry and confocal imaging

Adult PKD-tdTomato mice were first (30 minutes prior the procedure) injected with the non-steroidal anti-inflammatory Metacam (5mg/kg) and subsequently anesthetized with intraperitoneal administration of 100 mg/kg ketamine (Carros, France) and 10 mg/kg xylazine (Puteau, France). Before skin incisions, the local antalgic Lurocaine (5mg/kg) was injected subcutaneously. Animals were then transcardially perfused with 0.1 M PBS followed by 4% paraformaldehyde (PFA; Sigma-Aldrich) and tissues were immediately removed, post-fixed for 1 h in PBS 4% PFA at room temperature then transferred in PBS. The lumbar region of spinal cord was embedded in low-melting point agarose (4% solution, in PBS) and sectioned into 40 µm thick coronal sections with a vibrating microtome (Leica VT1000S). For KCC2 staining, free-floating sections were permeabilized for 1 h at room temperature using PBS with 0.2% Triton X100 (Sigma) and 3% BSA (bovine serum albumin). Sections were then blocked for 45 minutes at room temperature with PBS containing 1% NGS (normal goat serum) and 3% BSA followed by overnight incubation at room temperature with rabbit anti-KCC2 IgG (1:1000 dilution; #07-432, Merck-Millipore) (Yassin *et al*., 2014). For NKCC1 staining, sections were incubated for 45 minutes at room temperature with PBS containing 0.3% Triton X100, 1% BSA and 3% NDS (normal donkey serum) followed by a treatment with 1% SDS for 5 minutes. Tissues were then incubated overnight at 4°C with mouse anti-NKCC1 IgG (1:100 dilution; T4, Developmental Studies Hybridoma Bank) (Lytle et *al*., 1995). After primary antibody incubation, tissues were washed with PBS and incubated for 2 h with secondary antibodies conjugated to AlexaFluor 488 (1:400; Invitrogen). Sections were then rinsed with PBS, counterstained with DAPI (1:1000; Invitrogen) and mounted on gelatine-coated slides and coverslipped with home-made mowiol mounting medium for fluorescence microscope preparation. Confocal images were captured with a Zeiss LSM 700 laser scanning microscope and then processed with ZEN 2009 light Edition (Zeiss) and ImageJ 1.53s software (NIH).

### Acute spinal cord slice preparation

Coronal spinal cord slices were prepared as previously described (Gerstmann *et al*., 2022). Briefly, adult wildtype or transgenic mice were anesthetized with the ketamine-xylazine mixture and perfused intracardially with an ice-cold (0-4°C), oxygenated (95% O_2_/5% CO_2_) and low calcium/high magnesium cutting solution containing (in mM): NaCl 75, NaHCO_3_ 33, NaH_2_PO_4_ 1.25, KCl 3, CaCl_2_ 0.5, MgSO_4_ 7, glucose 15, sucrose 58, ascorbic acid 2, Na-pyruvate 2, *myo*-Inositol 3 (pH 7.3-7.4 and osmolality of 300-310 mosmole.kg^-1^). Following laminectomy and spinal cord dissection, lumbar spinal cord slices (250 µm thick) were cut with a vibratome (Leica VT1000S) in ice-cold cutting solution saturated with 95% O_2_-5% CO_2_. Slices were subsequently transferred to a holding chamber and incubated at 35° C for 15-20 minutes in oxygenated artificial CSF (aCSF) containing (in mM): NaCl 115, NaHCO_3_ 26, NaH_2_PO_4_ 1.25, KCl 3, CaCl_2_ 2, MgSO_4_ 2, glucose 15, ascorbic acid 2, Na-pyruvate 2, *myo*-Inositol 3 (pH 7.3-7.4 and osmolality of 300-310 mosmole.kg^-1^) and then kept at room temperature before recordings.

### Electrophysiology

Slices were placed in a recording chamber and superfused with oxygenated aCSF (1.5-2.0 mL/min) maintained at 30°C by an in-line peltier heater (Scientifica). Neurones around the central canal were visualized under infrared illumination with a 40x objective and oblique optics mounted on a Scientifica SliceScope Pro 1000. Data were acquired with an Axopatch 200A patch-clamp amplifier (Molecular Devices Inc.), low-pass filtered at 2 KHz and digitized at 10-20 KHz using a Digidata 1322A interface driven by pClamp 10 software (Molecular Devices). Patch pipettes (4-6 MΩ) were pulled from borosilicate glass capillaries (Harvard Apparatus) using the vertical PC-100 puller (Narishige International Ltd) and filled with an internal solution supplemented with 10 μM of the fluorescent dye AlexaFluor 594 (Invitrogen). In all experiments, CSF-cNs identity was confirmed by the recording of PKD2L1 channel activity as well as the visualisation of the dendritic protrusion (or bud) after whole-cell dialysis with Alexa 594.

#### GABA reversal potentials measurement

GABA_A_ current reversal potentials (E_GABA_) were determined with whole-cell voltage-clamp recordings using electrodes filled with a solution containing (in mM): K-Gluconate 119, KCl 21, NaCl 4, HEPES 5, MgCl_2_ 2, EGTA 1.1, Na_2_-ATP 2, Creatine-Phosphate 5, Na_3_GTP 0.6 (adjusted to pH 7.35 with KOH). Currents were evoked by brief (30 ms) puffs of GABA (1 mM) from a patch pipette at intervals of 10 s between applications and in the presence of DNQX (20 μM) and strychnine (1 μM) to block AMPA/kainate glutamatergic and glycinergic synaptic transmission, respectively. GABA I-V curves were obtained by averaging three consecutive responses for each voltage step and plotting the peak current amplitude as a function of the membrane potential. Experimental data points were fitted with linear regression and E_GABA_ was determined as the x-intercept of the linear fit. Membrane potentials were corrected offline for liquid junction potential (10 mV). Theoretical E_GABA_ was calculated with the Hodgkin-Katz-Goldman equation assuming a permeability ratio between Cl^-^ and HCO_3-_anions of 0.25 (Bormann *et al*., 1987; Kaila, 1994).

#### Cell-attached recordings

CSF-cNs membrane potential (V_m_) were estimated by measuring the reversal potential of K^+^ currents in cell-attached patches, as described previously (Verheugen *et al*., 1999; Fricker *et al*., 1999) with patch pipettes filled with a high K^+^ solution containing (in mM): K-Gluconate 140, NaCl 4, HEPES 5, MgCl_2_ 2, EGTA 1.1, Na_2_-ATP 2, Creatine-Phosphate 5, Na_3_GTP 0.6 (adjusted to pH 7.35 with 4 mM KOH). In these conditions, the [K^+^] in the pipette solution is close to the estimated intracellular [K^+^] in other cell types (Hille, 1992) so that the equilibrium potential for K^+^ (E_K_) across the patch would be around 0 mV and K^+^ currents reverse when the pipette potential (V_pip_) cancels V_m_ out. Therefore, the neuron holding potential (−V_pip_) at which K^+^ currents reverse gives a direct and non-invasive quantitative measure of V_m_ (at K^+^ reversal, V_patch_ = V_m_-V_pip_ ≈ 0 mV). Voltage-gated K^+^ channels were activated with voltage ramps (−140 to +200 mV for 100 ms) applied every 1 s, in control and during pressure application of GABA (1 mM, 10 s duration). K^+^ currents were recorded in the presence of TTX (0.5 μM), DNQX (20 μM) and strychnine (1 μM). E_K_ was measured from the intersection between the K^+^ current and the fit to the linear component.

The effect of GABA on CSF-cNs firing activity was investigated with tight-seal cell-attached recordings in voltage-clamp mode by adjusting the command potential to result in a holding current of 0 pA (Perkins, 2006). Action potential (AP) currents were recorded in control and during pressure application of GABA (1mM, 5 s duration). In each cell and before AP recording, the effect of GABA on CSF-cNs membrane potential (depolarising or hyperpolarising) was assessed with cell-attached current-clamp recordings in tight-seal configuration and with pressure application of the agonist (1 mM, 5 s duration; see Results). Voltage-clamp and current-clamp cell-attached recording were performed with patch pipettes filled with K-gluconate based internal solution.

#### Recording of inwardly-rectifying (GIRK/kir3) potassium currents

Postsynaptic GIRK-type K^+^ currents were recorded in voltage-clamp whole-cell configuration with patch electrodes filled with a solution containing (in mM): 140 K-gluconate, 4 NaCl, 5 HEPES, 2 MgCl_2_, 1.1 EGTA, 2 Na_2_ATP, 5 Na_2_-phosphocreatine, 0.6 Na_3_GTP (adjusted to pH 7.35 with KOH). GIRK currents were evoked by bath application of GABA_B_-Rs selective agonist baclofen (100 μM) or the potent activator of GIRK1/2 channels ML297 (VU0456810, 100 μM) (Kaufmann *et al*., 2013) at a holding potential of -50 mV and in the presence of antagonists DNQX (10 μM) and strychnine (1 μM).

### Calcium imaging

CSF-cNs selectively expression the fluorescent probe GCaMP6f were illuminated by light through an 490/30 nm (peak/bandwidth) excitation filter using the CoolLED epifluorescence system (p1 PrecisExcite). Emitted light was bandpass filtered at 535/30 nm and collected with the HQ2 CoolSnap CCD camera (standard mode at 10 MHz and Gain 1, Photometrics) connected to a PC through a frame grabber (CoolSNAP LVDS interface card, Photometrics) and controlled by the MetaView software (Molecular Devices Inc.). Images were acquired in time lapse mode with a 2×2 binning for 100 ms at 2 Hz for 70 s period (141 images). For each image, a region of interest (ROI) was drawn over the CSF-cNs soma as well as the background and the average intensity of the fluorescence in these two ROIs was calculated. Subsequently, the background fluorescence intensity was subtracted from the somatic data to compute the net fluorescence (F) and data were expressed as the change of fluorescence relative to basal fluorescence (F-F_*0*_/F_*0*_ or ΔF/F_*0*_). Modifications in intracellular Ca^2+^ concentration ([Ca^2+^]_i_) were evoked at room temperature by pressure application of GABA (1 mM, 10 s duration) or high-potassium aCSF (20 mM, 10 s duration) in the presence of DNQX (20 μM), strychnine (1 μM) and the selective antagonist CGP54626 (10 μM) of GABA_B_-Rs to prevent their inhibitory effect on voltage-gated calcium channels (Jurčić *et al*., 2019). In experiments with calcium-free aCSF (0-Ca^2+^), slices were perfused with external solution containing (in mM): NaCl 115, NaHCO_3_ 26, NaH_2_PO_4_ 1.25, KCl 3, MgSO_4_ 4, glucose 15, ascorbic acid 2, Na-pyruvate 2, *myo*-Inositol 3, EGTA 0.2.

### Data analysis and statistics

Electrophysiological data were analysed using Clampfit 10 (Molecular Devices Inc.), Excel 2016 (Microsoft) and Prism 9 software (GraphPad). Data were expressed as mean ± SD in text and represented as box-whisker plot using the Tukey’s method with Prism 9 software. The box plot represents the distribution of data as a box, with the median as a central line and the hinges as the edges of the box (lower and upper hinges corresponding to the 25^th^ and 75^th^ percentiles, respectively). The upper whisker extends to the largest value in the data set that does not exceed 1.5 times the interquartile range (IQR) plus the 75^th^ percentile (where IQR is the difference between the 25^th^ and the 75^th^ percentiles). The lower whisker extends to the smallest value higher than the 25^th^ percentile minus 1.5 times IQR. Data beyond the end of the whiskers are called outliers and are plotted individually as circles. For all electrophysiological data and histological experiments, “n” refers to the number of recorded cells and to the number of mice exanimated, respectively. All data were tested for normal distribution using the Shapiro-Wilk test with Prism 9 and the result determined the use of parametric or nonparametric statistical tests indicated in the Results section. Fisher’s exact test was used to compare 2 × 2 contingency tables. Statistical tests were performed with Prism 9 and differences were considered significant when p < 0.05.

### Reagents

All reagents were purchased from Sigma-Aldrich except otherwise stated. Tetrodotoxin (TTX) was from Alomone Labs. 6,7-dinitroquinoxaline-2,3-dione disodium salt (DNQX) and ML297 (VU0456810) were from Abcam Biochemicals. (R)-baclofen, gabazine (SR 95531), CGP54626 and furosemide were from Tocris Bio-techne.

## Additional information

### Data availability statement

The data of this manuscript are available from the corresponding author upon reasonable request.

### Authors contributions

All experiments were performed in laboratories at the ‘Institut des Neurosciences de la Timone (INT)’ of Aix-Marseille University. Conception and design of the work: J.T., N.W. & R.S. Acquisition, analysis and interpretation of data for the work: P.R., N.J., N.W. & R.S. Drafting the work or revisiting it critically for important intellectual content: P.R., N.J., J.T., N.W. & R.S. All authors have read and approved the final version of this manuscript and agree to be accountable for all aspects of the work in ensuring that questions related to the accuracy or integrity of any part of the work are appropriately investigated and resolved. All persons designated as authors qualify for authorship, and all those who qualify for authorship are listed.

## Acknowledgments

We gratefully thank Caroline Blanc-Tailleur and Anne Kastner for her assistance in immunohistochemistry experiments. We acknowledge the ‘Institut de Neurosciences de la Timone (INT)’ technical facilities for their support in the study (Neuro-Bio-Tools: Molecular Biology and Histology; INPHIM: confocal microscopy).

